# A highly parallel strategy for storage of digital information in living cells

**DOI:** 10.1101/096792

**Authors:** Azat Akhmetov, Andrew D. Ellington, Edward M. Marcotte

**Author notes:** Correspondence to (AA), (EMM).

## Abstract

Encoding arbitrary digital information in DNA has attracted attention as a potential avenue for large scale and long term data storage. However, in order to enable DNA data storage technologies there needs to be improvements in data storage fidelity (tolerance to mutation), the facility of writing and reading the data (biases and systematic error arising from synthesis and sequencing), and overall scalability. To this end, we have developed and implemented an encoding scheme that is suitable for detecting and correcting errors that may arise during storage, writing, and reading, such as those arising from nucleotide substitutions, insertions, and deletions. We propose a scheme for parallelized long term storage of encoded sequences that relies on overlaps rather than the address blocks found in previously published work. Using computer simulations, we illustrate the encoding, sequencing, decoding, and recovery of encoded information, ultimately demonstrating the possibility of a successful round-trip read/write. These demonstrations show that in theory a precise control over error tolerance is possible. Even after simulated degradation of DNA, recovery of original data is possible owing to the error correction capabilities built into the encoding strategy. A secondary advantage of our method is that the statistical characteristics (such as repetitiveness and GC-composition) of encoded sequences can also be tailored without sacrificing the overall ability to store large amounts of data. Finally, the combination of the overlap-based partitioning of data with the LZMA compression that is integral to encoding means that the entire sequence must be present for successful decoding. This feature enables inordinately strong encryptions. As a potential application, an encrypted pathogen genome could be could be distributed and carried by cells without danger of being expressed, and could not even be read out in the absence of the entire DNA consortium.

## Introduction

Recently, the prospect of encoding information in free nucleic acids has attracted much interest from both academic research communities [1] [2] [3] as well as the technology sector [4]. DNA offers unique potential for storage of information, in that large amounts of information can be be written (synthesis) and read (sequencing) at moderate, and rapidly decreasing, cost. Ultimately, one DNA base-pair (bp) stores 2 bits of information [1], a much more dense information storage medium than any electronic device of comparable capacity. Moreover, the long term storage of information in DNA becomes easy and practical, given its extremely long half-life [5], unlike digital media which is prone to degradation. As an example, while it has been possible to reconstruct a mammoth genome from remains found in the tundra [6], it is unlikely we would recover electronic information stored in the same way. This feat was possible in part because an exquisite molecular mechanism (base-pairing and replication) exists for making many copies with very high fidelity.

Herein we propose a novel scheme for encoding information in DNA and distributing this information across multiple cells, and present computer simulations demonstrating the feasibility of our approach. We discuss three main steps of this process (Figure 1): (i) A coding scheme for converting digital information into DNA and vice versa. Our scheme has a built-in, highly general error detection and correction capacity that can be tailored to pre-chosen balances of redundancy versus error tolerance; (ii) A strategy for parceling long strings of information into smaller pieces that allows for their later re-assembly. This strategy is completely compatible with current chemical DNA synthesis methods that yield at most only short (~200 bp) oligonucleotides; and (iii) The use of error tolerance to defeat corruption of information arising from synthesis errors, sequencing errors, mutations and packet loss. Taken together, these innovations allow for very long term, error-resistant storage, potentially for thousands of years.

**Figure 1:**
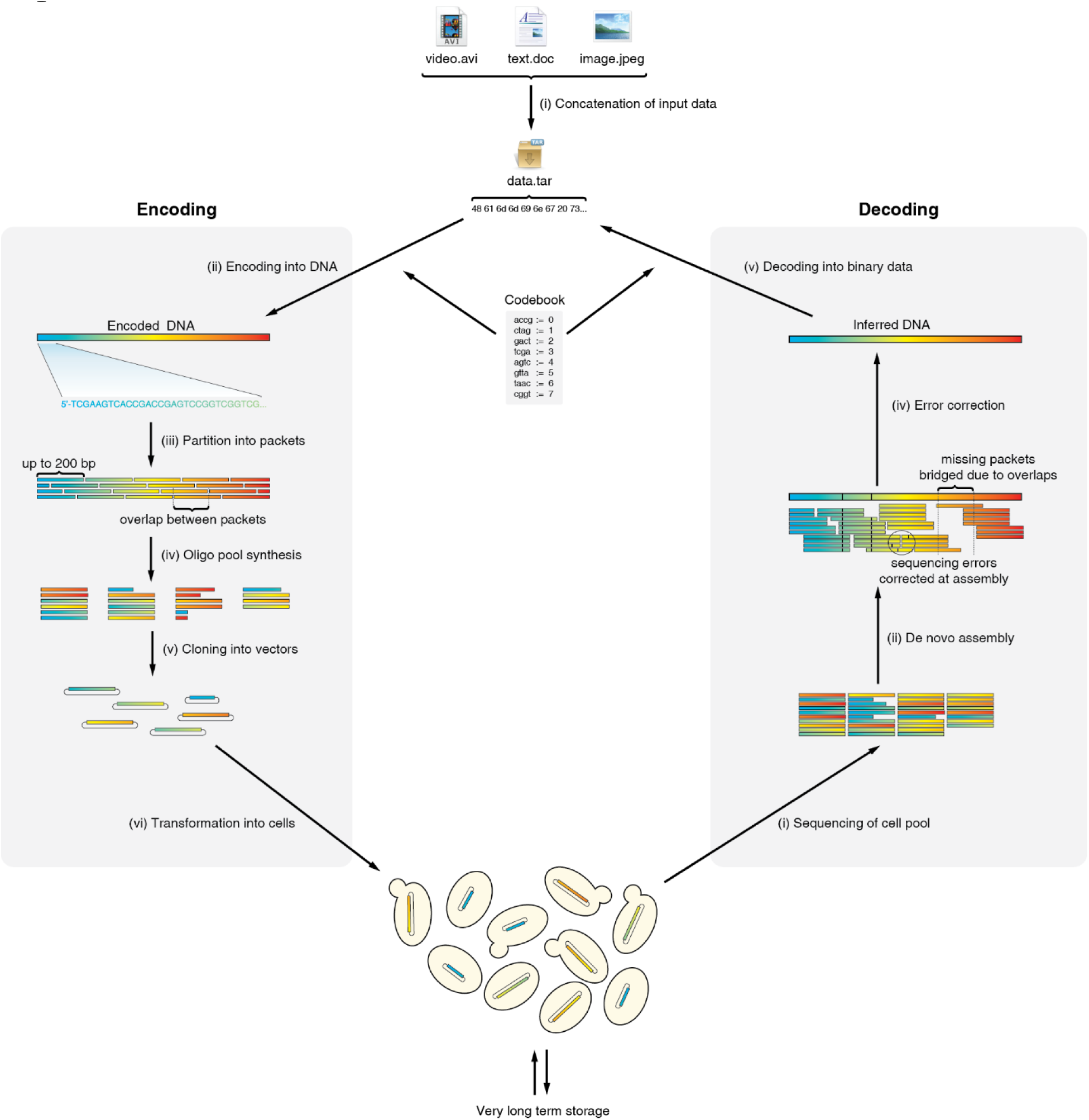
A diagram of the encoding and decoding process. The input data is first wrapped in a tar archive, to ensure a uniform input format as well as combining multiple files into a single contiguous data stream with a well-established method. The digital information is encoded using a pre-generated codebook, producing one long single sequence of DNA. This sequence is split into overlapping packets, each up to 200 bp long, which are then synthesized as a complex pool of oligonucleotides. These can be cloned into plasmids and transformed into cells, where they can be maintained reliably for a very long time. To recover the information, the population of cells (or alternatively plasmids or lyophilized oligonucleotides) can be sequenced with NextGen sequencing technology, and *de novo* assembly of the resulting reads is performed. During assembly, some errors can be corrected by simply considering the consensus of the contig, whereas systematic errors (such as those arising during synthesis) can be corrected *in silico* using the error correcting code. Finally, the codebook is used to decode the resulting contig and recover the digital files.

We also analyze the performance of our method for key trade-offs that will be inherent in any strategy of encoding digital information in DNA.

## Methods

### Test data

The Hamming image was obtained by scaling down a color photo of Richard Hamming (available under the Creative Commons license) and encoding with the Jpeg algorithm. After adjusting the pixel size of the image to approximate our desired size, we then fine-tuned the Jpeg compression level (ultimately we used level 86) so that the resulting file size was about 10 kB. The random data file was obtained by creating a 10 kB file and populating it with random data piped from /dev/urandom on a Unix computer. The centromere input was a text file containing part of the sequence from human chromosome I (NCBI sequence NC_000001.11 positions 11000-22000). The flat file was constructed by pasting a large number of 0s into a text file. In every case, the test data was wrapped in a tar archive (which is a file format that can store one or more files without compression), which is then used as the input for the DNA codec.

### Codebook generation

The codebook is a table showing permissible blocks of DNA sequence, and what numeric value each sequence maps to. Our algorithm (implemented as Python code in the file make_codewords.py) begins by generating all possible sequences of a given length (Table 1). This pool is filtered to remove sequences with high repetitiveness (quantified by dividing the number of its unique subsequences to the number of all of its subsequences, implemented in complexity_estimation.py) and undesirable GC content (in our case set as less than 40% or more than 60% GC). Of the remaining sequences, one is picked at random and saved as a codeword. All sequences with Levenshtein distance less than the defined threshold (in our case 3) are removed. Of the remaining sequences, another one is picked at random to be the second codeword, those with too small Levenshtein distance are pruned again, and the process is repeated until no further codewords can be produced. Sequential integer values starting from 0 are then assigned at random to each codeword in the resulting codebook. We generated a single codebook of eight 4 bp codewords and used it for all of our experiments (Table 2).

**Table 1:** Parameters used for codebook generation

| Parameter | Explanation | Value |
| --- | --- | --- |
| Word length | Length of each codeword to be generated, also determines cipher block size | 4 |
| Max mutations | Maximum number of mutations that can be recovered from per block. If this parameter is $k$, the algorithm generates codewords such that the minimum Levenshtein distance between them is $2k + 1$. | 1 |
| Min. GC | Minimum GC composition in each codeword | 0.4 |
| Max. GC | GC composition in each codeword | 0.6 |
| Complexity | Minimum complexity (used as a proxy for repetitiveness) of each codeword | 0.75 |

**Table 2:** Codebook

| Codeword | Value |
| --- | --- |
| accg | 0 |
| ctag | 1 |
| gact | 2 |
| tcga | 3 |
| agtc | 4 |
| gtta | 5 |
| taac | 6 |
| cggt | 7 |

### DNA codec implementation

The DNA encoding-decoding scheme was implemented in Python (dna_read.py and dna_write.py) to produce a codec which, given the codebook, converts a digital file into a DNA sequence (encoding) and vice versa (decoding). For encoding, the file is first read, compressed with LZMA (lzma library of Python 3.5.2), converted into a byte array, which is equivalent to the digits of a base 256 number. A base change operation is performed, to change the base from 256 (the number of possible values a byte may have) to 8 (the number of codewords in our codebook). The resulting array of integers is converted into short DNA blocks according to the mapping in the codebook, and concatenated into a single DNA sequence. To decode, the entire DNA sequence is broken into 4 bp blocks, then each block is mapped back to an integer according to the codebook, producing a base 8 number. A base change is performed from 8 to 256, and the resulting array of numbers is converted into a byte stream and decompressed with LZMA.

### Error correction

The error correction is applied by scanning through a DNA sequence and checking each subsequent block against the codebook. If the block appears in the codebook, then it is deemed correct. Otherwise, the block is deemed to be mutated, and the Levenshtein distance from the observed block to each code word is calculated. The block with the minimum distance is assumed to be the original sequence before mutation. This was implemented in clean_dna.py and related files.

### Partition of encoded DNA into packets

In order to accommodate eventual synthesis of the encoded DNA sequence into nucleic acid, it is broken up into short subsequences termed “packets” (implemented in split_into_packets.py). The packets are selected so that each one is 200 bp long (selected because it is the longest length feasible with current largescale DNA synthesis technology), and they tile original sequence such that each two adjacent packets overlap by a specified amount. For the beginning and end of the sequence, shorter packets are produced to avoid lower coverage of sequence ends.

### Sequencing simulations

We simulated NextGen sequencing of our encoded DNA and packets using the ART sequencing simulator (Huang 2012). We simulated 150 bp-long reads with the Illumina HiSeq 2500, this being a common, commercially available method of sequencing. The read depth varied from 1x to 50x depending on the particular experiment.

### *De novo* assembly

Fastq files generated by the ART simulator were used for *de novo* contig assembly with the Geneious assembler from the Geneious 9.1.2 software [7]. The parameters specified that the expected assembly was linear and not circular. The longest resulting contig was then taken as the assembled sequence.

### Mutation simulation

To simulate mutations, we implemented a simple iterative algorithm (mutate_dna.py). In each iteration, the algorithm picks a random base pair and changes it to a different, randomly picked base. This operation is repeated as many times as needed.

## Results

### Successful generation of codebook

We executed our implementation of the codebook generation algorithm to generate a set of keywords 4 bp long, and with minimum Levenshtein distance [8] of 3 (the full set of parameters is given in Table 1: Parameters used for codebook generation). The latter parameter was set so as to allow recovery from up to one mutation (including substitutions, insertions and deletions) per block of encoded DNA. Due to a codeword length of 4, the block length is likewise 4 bp, therefore the expected theoretical upper bound on mutation rate for error recovery is 0.25 bp^−1^, and the upper bound for error detection is 0.5 bp^−1^.

All of the resulting code words showed a good diversity of base pair composition, lack of repetition, sufficient sequence distance between themselves, and overall conformed to expectations. The list of keywords is given in Table 2 along with the numeric value assigned to each codeword for the work described here.

The theoretical upper bound on the information that can be stored using the four nucleotides of DNA is 2 bits bp^−1^. Given our codebook, each sequence of 4 bp can only have one of 8 values, therefore the information content under our encoding scheme is only 3 bits per 4 bp, with a density of 0.75 bits/bp. Thus, the expected theoretical rate of our encoding approach *per se* can be calculated as 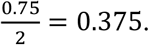

### Encoding of digital data into DNA

We implemented our encoding algorithm and used it to encode four separate sets of input data (Table 3). Table 4 describes the performance of our encoding algorithm on these test data sets. Each input file was wrapped in a tar archive prior to encoding, and rates were calculated by dividing the size of the tar file prior to converting (found by multiplying the size in bytes by 8) by the information content of the resulting DNA (found by multiplying the number of base pairs by 2).

**Table 3:** Input data

| Input data | Explanation | Rationale |
| --- | --- | --- |
| Hamming.jpeg | Color photo of Hamming in jpeg format, scaled down to 161x237 pixels | Example of real world data |
| Flat file | A text file containing a string of 10,340 zeros | Demonstrate performance when given extremely repetitive data |
| Random data | 10 kb of random data obtained from /dev/urandom on a Linux computer | Demonstrate performance when given data without any statistical bias |
| Centromere | Part of centromeric sequence from human chromosome I, retrieved from:<br>https://www.ncbi.nlm.nih.gov/nuccore/NC_000001.11?report=fasta&log\$=seqview&format=text&from=11000&to=22000 | Real-world example of repetitive information |

**Table 4:** Data before and after compression

| **Data** | **Size of file** | **Size of tar archive** | **Length of encoded DNA** | **Rate** |
| --- | --- | --- | --- | --- |
| Hamming.jpeg | 10,478 | 12,288 | 111,192 | 0.442 |
| Flat file | 10,340 | 11,776 | 1,792 | 26.286 |
| Random data | 10,240 | 12,288 | 112,044 | 0.439 |
| Centromere | 11,001 | 12,800 | 35,200 | 1.454 |

#### Hamming image

The realistic test input Hamming.jpeg (Figure 2) was encoded and had an empirical rate of 0.442, thus the redundancy and overhead introduced by our encoding algorithm inflated the size of the data by 1.262 times. Notably, this is a rate higher than the predicted 0.375; the most likely explanation for this is that the LZMA compression step built into our method was successful in reducing the size of the input. After decoding, we were able to recover the original image exactly.

**Figure 2:**
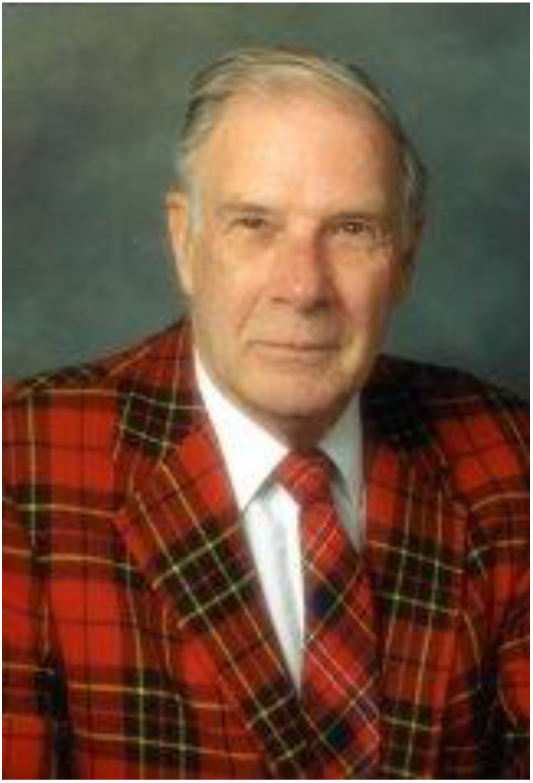
Digital data used for *in silico* experiments. Left: A 161×237 pixel color photo of Richard Hamming, encoded with the Jpeg algorithm so as to produce a file 10,478 byte file.

The resulting DNA string appeared to be free of any major self-similarity or repetition. We visualized the self-similarity using a dot plot to show the extent to which the sequence matches itself (Figure 3). The most notable case of repetition was near the beginning and ends of the sequence (Figure 4). Closer inspection of the binary data after compression but prior to transformation into DNA revealed that there is often a short string of zeros in LZMA-compressed data, containing header/tail information used by the compression logic to identify the properties of a compressed data stream. Indeed, the most commonly repeated sequence at these regions was ACCG, which maps to 0 in our codebook.

**Figure 3:**
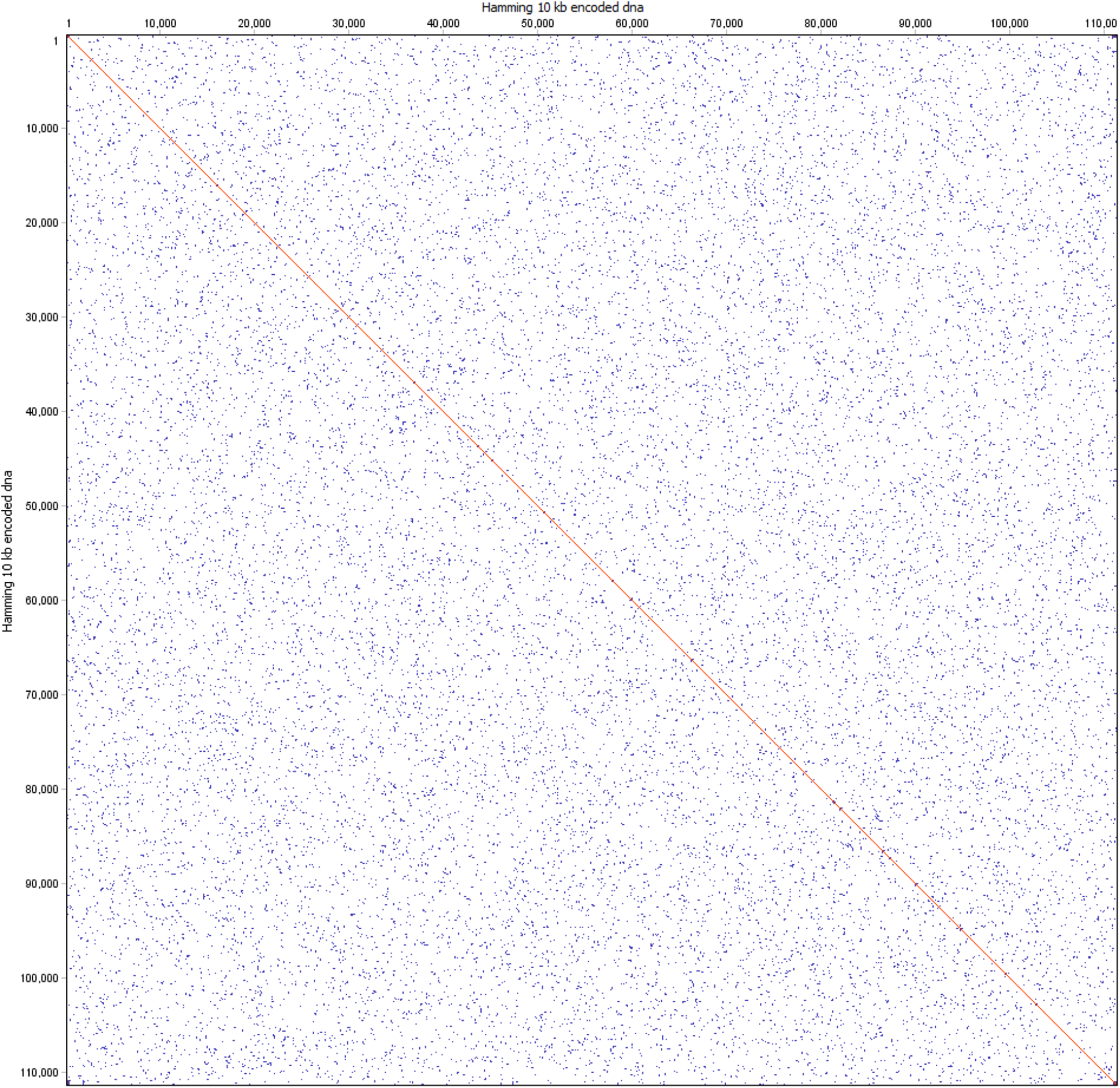
Overall self-similarity of the encoded Hamming image. Dot plot of the encoded Hamming image generated with dottup, using word size 20 as the parameter. Positions where 20 bp of the sequence are self-similar are marked with blue. Identical regions longer than 100 bp are marked with red. The plot shows a lack of long stretches of repetition that could interfere with assembly.

**Figure 4:**
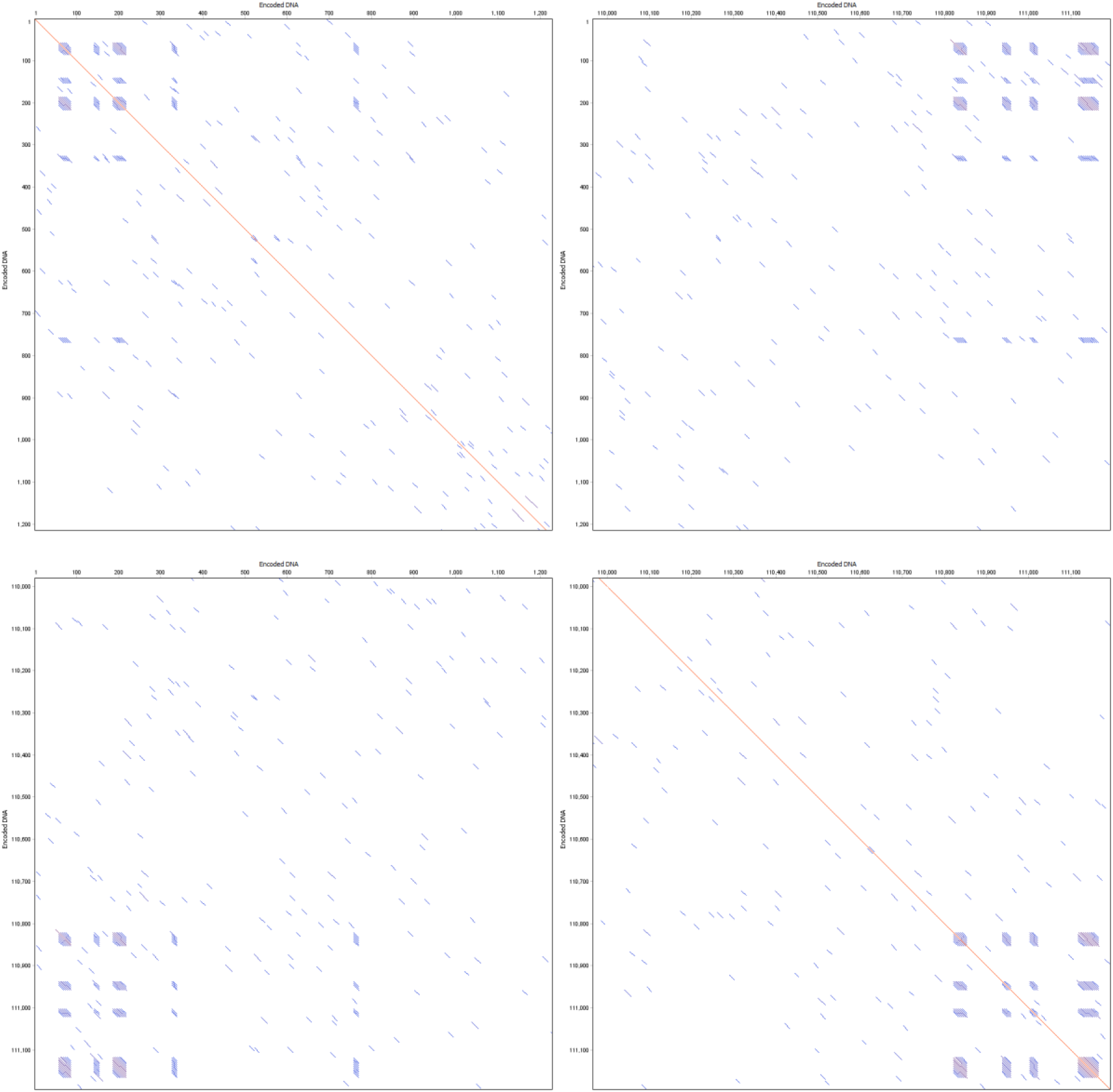
Self-similarity at corners. Dotplots of the same encoded DNA, showing only the ends of the sequence, generated with word size 10. Short blocks of repetitive sequence are visible as blue blocks, these result from header and terminator information utilized by the LZMA algorithm which is less variable than the compressed data stream itself. Top left: Sequence head vs. itself. Top right and bottom left: Head vs. tail. Bottom right: Sequence tail vs. itself.

#### Random data, centromere and flat file

Random data is often regarded as particularly difficult to compress, due to a lack of statistical tendencies in it which can be exploited by compression strategies. Indeed, upon compressing our 10 kb random file with 7zip, an open source implementation of the LZMA algorithm, we obtained a compressed file 10.2 kb in size. When encoded into DNA with our algorithm, the resulting file was only 1.278 times bigger than the input tar file.

The centromere was included as an example of highly repetitive information that has statistical properties representative of known repetitive DNA that is considered challenging to sequence and synthesize. As seen by the high rate, the compression step of our encoding process vastly reduced the size needed to store this data. The resulting DNA string had very similar composition and structure to the other test data, and should be no more difficult to synthesize or read with our approach than any other input.

Lastly, the flat file was dramatically reduced in size after compression, as shown by the extremely high rate. Due to the very short output sequence, the repetitive head/tail regions are clearly visible in the dot plot (Figure 5), as the data content of the end result is relatively small.

**Figure 5:**
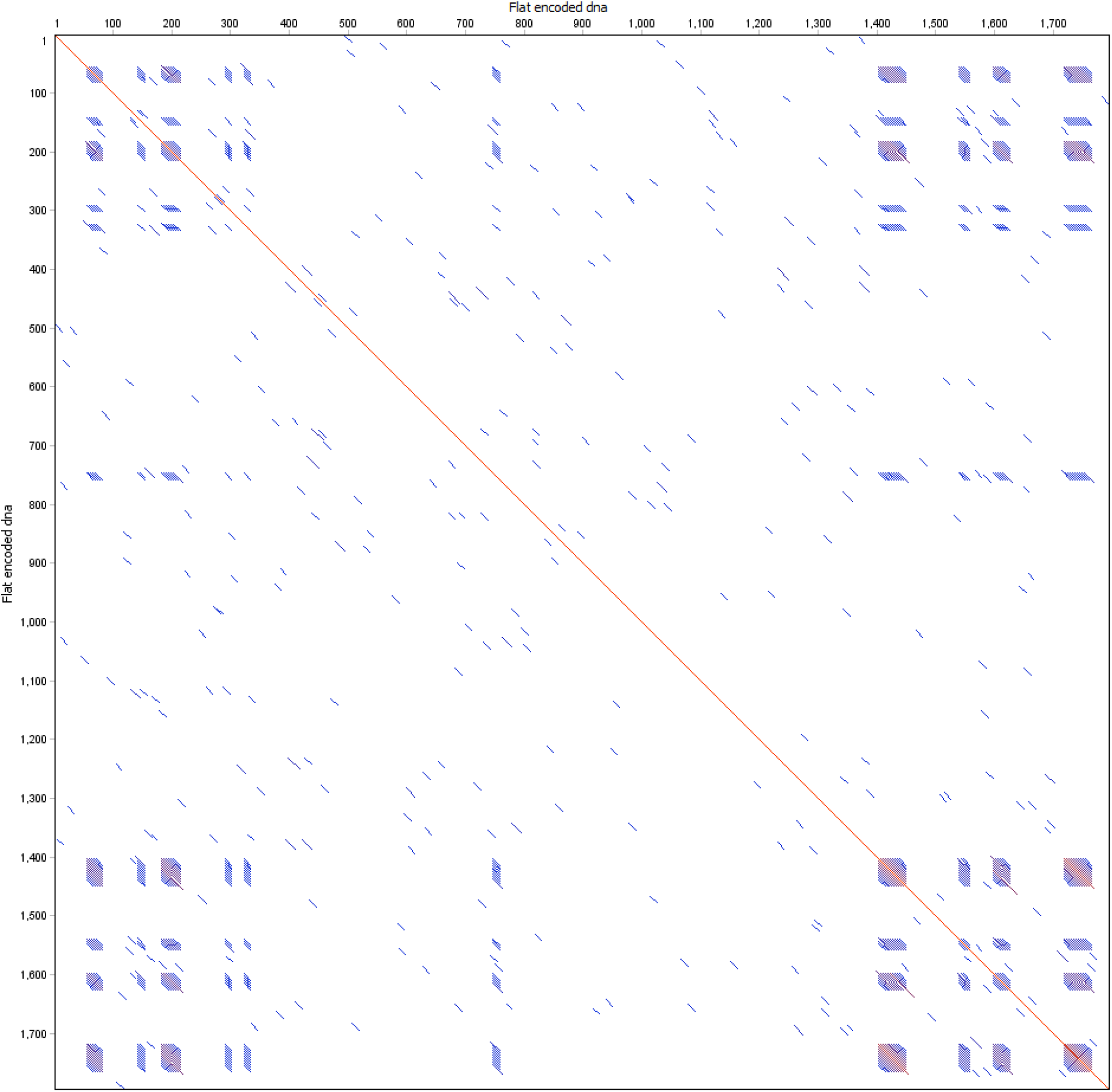
Self similarity of flat file. Dot plot of the entire encoded flat file, generated with dottup with word size 10, showing self-similarity within the entire encoded DNA sequence.

#### Base pair composition

We noted that the nucleotide composition was very close to an even split of 25% for each base in most of our encoded DNA (Figure 6). The most conspicuous exception was the flat file, which showed larger skews due to the characteristic head and tail patterns, which do not vary in length with the amount of data encoded, thus having a disproportionate effect.

**Figure 6:**
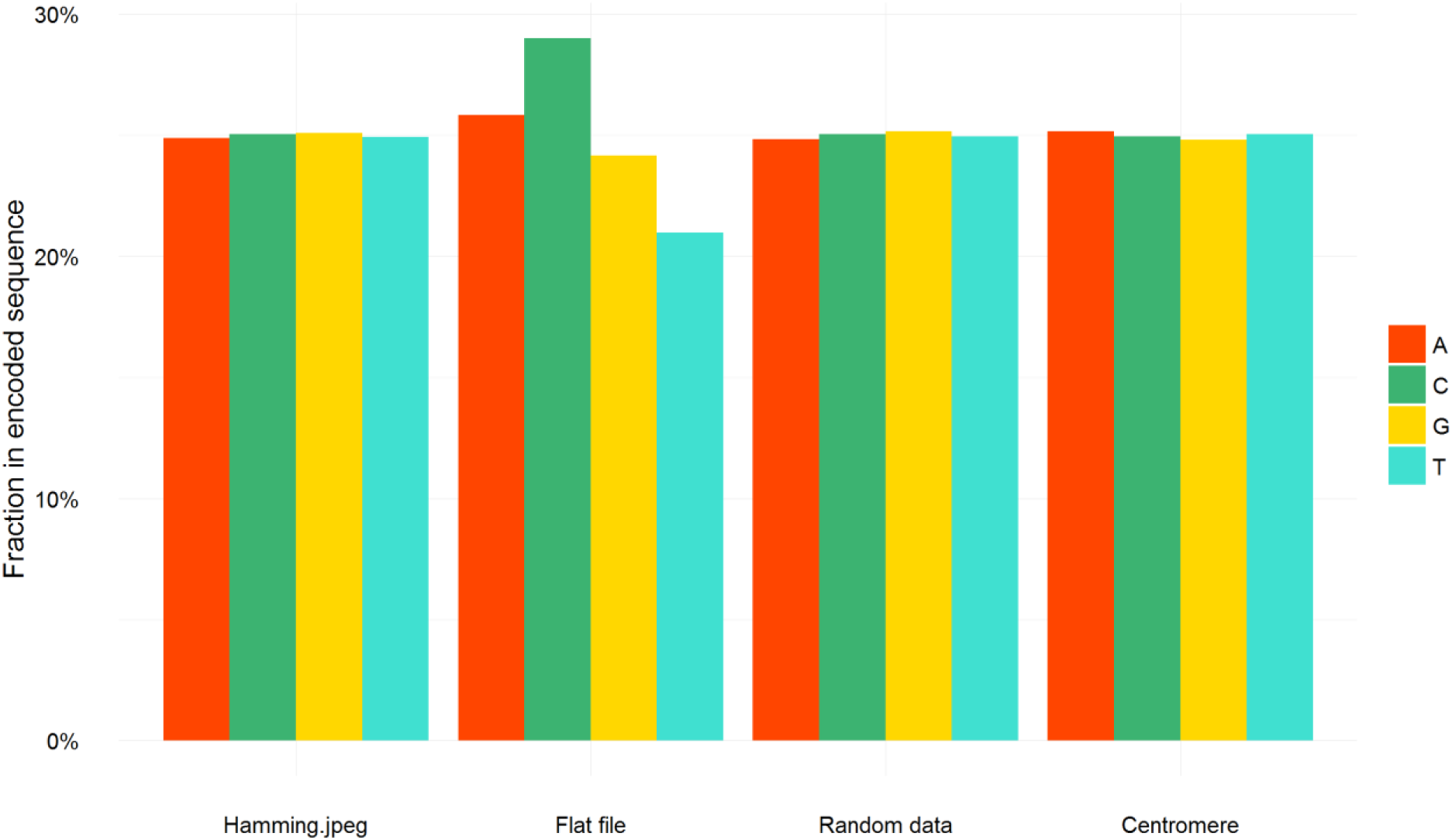
Total nucleotide composition of encoded DNA. Bars show the relative fraction of each nucleotide within DNA obtained by encoding the given digital data.

We have also investigated the local composition to detect any small stretches of base composition skew (Figure 7). As expected, we did not observe any regions of pronounced skew, and the distribution of nucleotides seemed to be uniform throughout the encoded sequences. Notably the skew caused by characteristic head and tail patterns is most noticeable in the flat file, where several peaks of C content can be seen at the beginning and end. This would be the expected result, since the common code word in these areas is ACCG which has a larger proportion of Cs. After quantifying the total skew by adding up local deviations from expected even distribution, we observed mostly small amounts of error, with the exception of the flat file which had larger composition imbalances, particularly in C and T.

**Figure 7:**
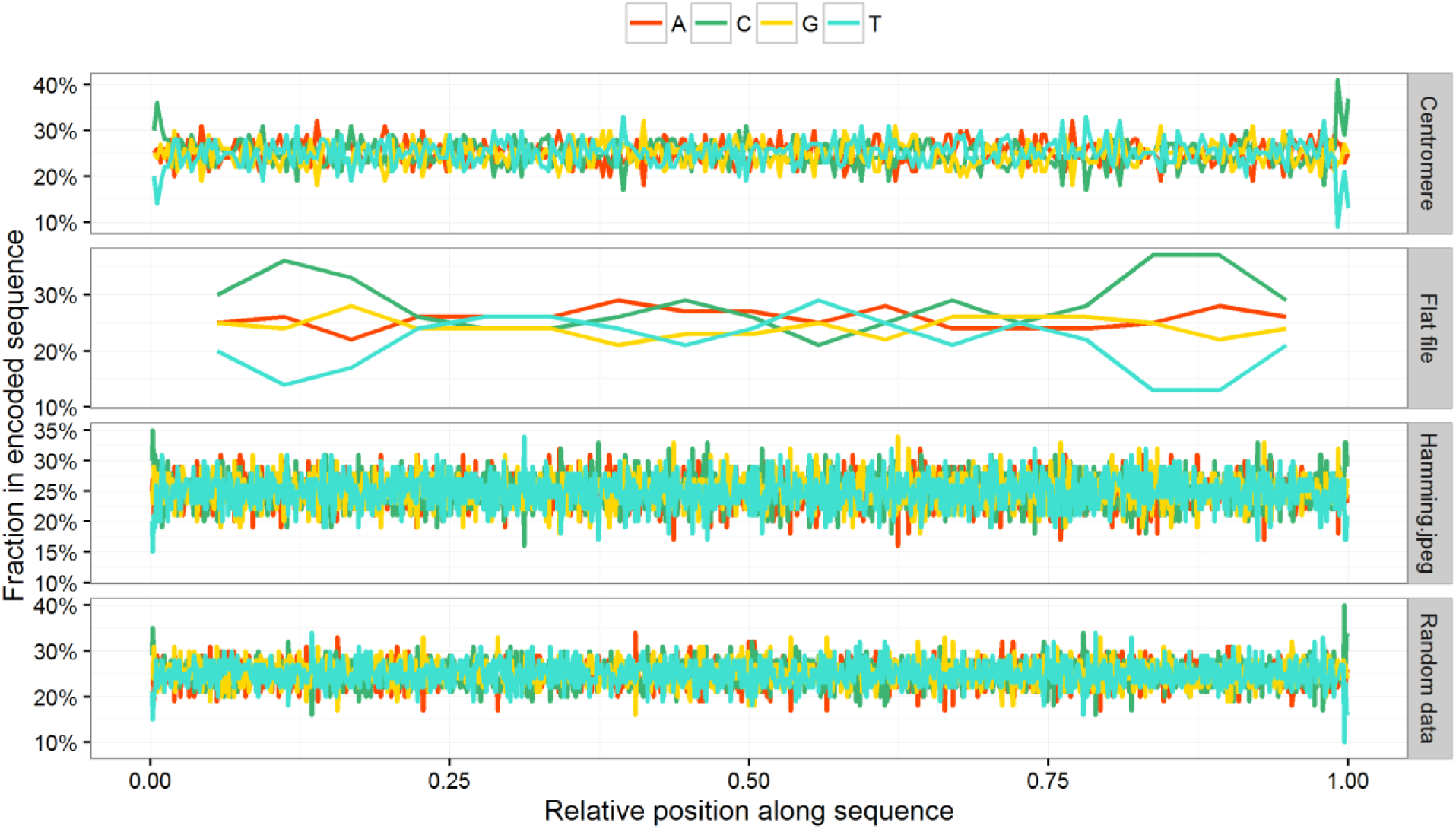
Local composition. Nucleotide composition in sliding 100 bp window for each sequence of encoded DNA.

**Figure 8:**
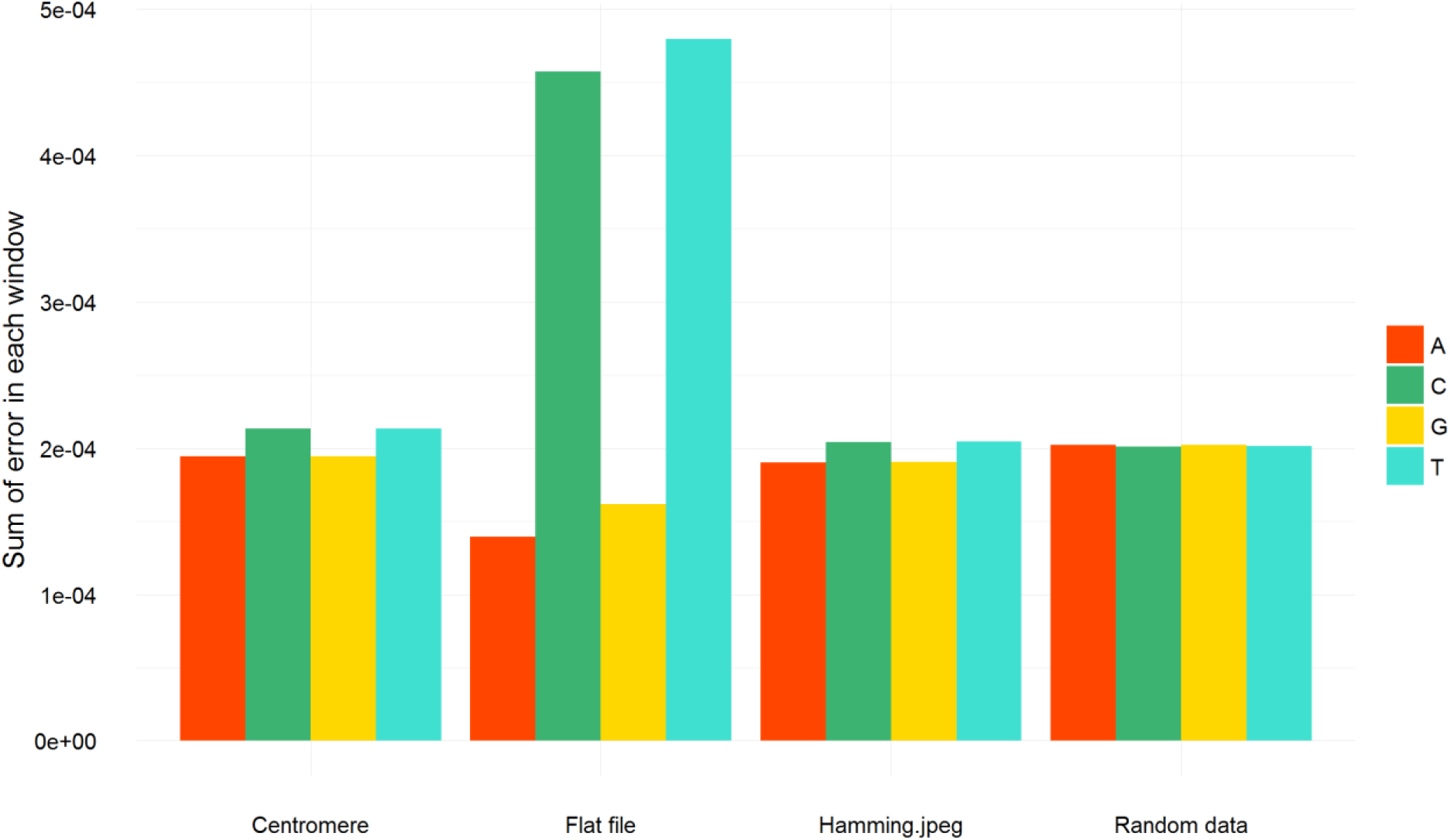
Total nucleotide composition error. Total deviation of nucleotide composition from the expected 25% proportion. Shown here is sum of error within each 100 bp window tiled along the encoded sequence, and divided by the sequence length.

#### Open reading frames within the encoded DNA

Encoded DNA produced by our algorithm is expected to resemble random sequence in many respects. It is possible that large stretches of such sequence would coincidentally contain start and stop codons forming open reading frames. We analyzed the occurrence of such random ORFs, and a visualization is presented in Figure 9.

**Figure 9:**
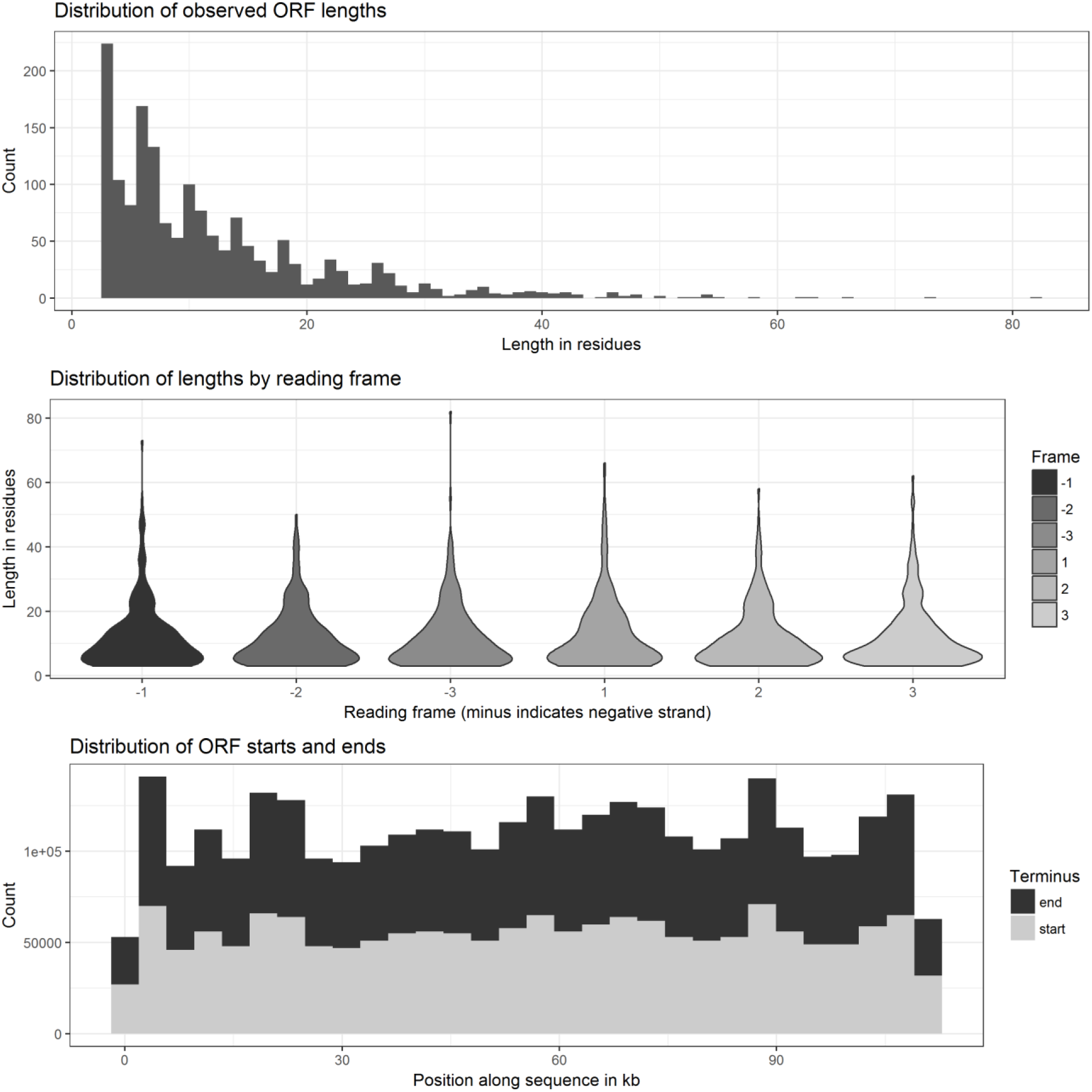
Spurious ORFs in encoded sequence. Top: Histogram showing the distribution of spurious ORFs observed in the DNA sequence for the encoded Hamming image. Middle: Violin plot showing the length distribution of spurious ORFs grouped by reading frame. Frames are marked with a minus (-) if they are on the negative strand (ie. detected in the reverse complement of the sequence). Bottom: Distribution of spurious ORF start (grey) and stop (black) positions along the sequence.

The distribution of ORF lengths follows a pattern that would be expected *a priori* from random sequence. The shortest ORFs are most frequent (our ORF detection software looks for ORFs containing at least one codon besides start and stop). ORFs longer than about 30 residues or 100 bp are very rare, and none are longer than 82 residues. The start and stop codons are evenly distributed throughout the sequence, but slightly under represented near the ends of the DNA – possibly owing to the composition skew introduced by the characteristic head and tail regions resulting from our encoding approach. There does not appear to be any particular bias towards a single reading frame, nor towards a particular direction.

### Decoding and error correction

We confirmed correct operation of our encoding approach by attempting to decode the DNA resulting from the encoding step. In every case we were able to recover the original tar archive, which when extracted produced the relevant file that was identical to the one encoded.

Having verified successful round-trip encoding-decoding of information we then sought to measure the performance of the error correction function. We simulated substitution mutations accumulating at a slow but steady rate over many generations, by repeatedly mutating the Hamming.jpg so as to correspond to total numbers of mutations per block ranging from 0 to 2. As an indicator of data integrity, we compared sequence identity between the simulated mutant sequence and the original, before and after applying the error correction (Figure 10). For a smaller number of mutations, the error correction is able to restore nearly the entire original sequence. As the density of mutations increased, eventually the error correction did become overwhelmed, but the data shows a very clear mutation buffering effect contributed by it.

**Figure 10:**
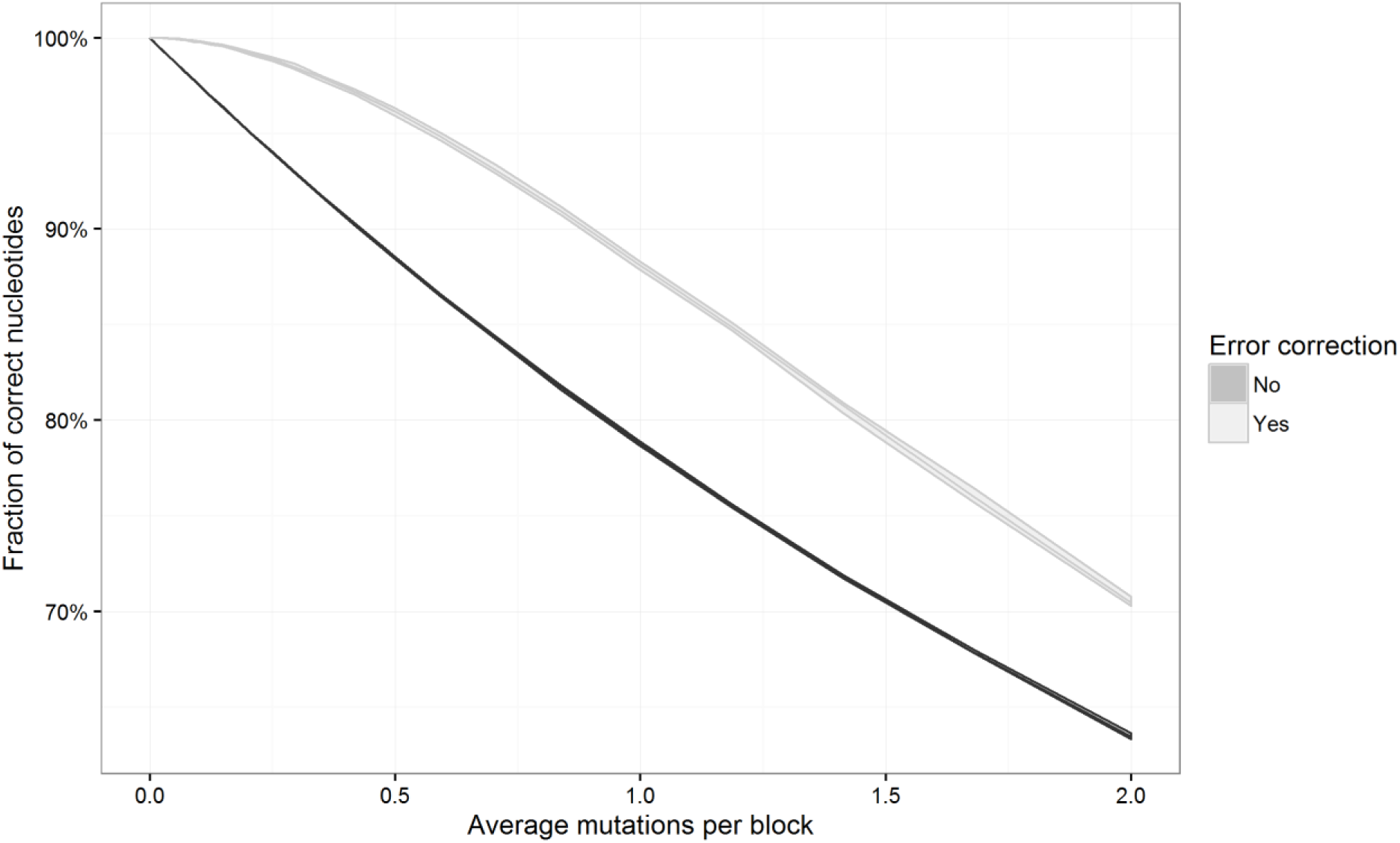
Mutation buffering by error correcting code. Light gray line shows the effect of applying error correction on sequences mutated to varying degrees. The mutation rate is shown in average mutations per block, given that each block is 4 bp long. Shown here is the mean of 10 simulations (middle line) with ±2.58σ band (shaded area), representing 99% confidence interval. Bottom plot: Subset of the data corresponding to only lower mutation rates.

As calculated in earlier sections, our error correcting code operates on a block-wise basis. Each individual block (in our case 4 bp long) can be recovered so long as no more than one mutation occurs within it. With larger numbers of mutations distributed randomly throughout the sequence, it becomes more probable that at least two mutations will fall very close to each other and coincide on the same block. Such an improbable event would potentially result in the loss of that block of information. We have observed that minor perturbations to the encoded DNA do not significantly corrupt the stored data, and do not preclude its recovery. Furthermore, our strategy anticipates that the final encoded DNA string will be broken up into small, overlapping pieces in practice; therefore the mutations would be further removed at the assembly stage via a consensus mechanism.

### Parallelized storage

We investigated the practical aspects of synthesis, storage and reading of digital data as DNA using our approach by simulating the round trip process. Basing our model on the assumption of DNA synthesis capability which allows the production of a pool of oligonucleotides 200 bp each (readily achievable with current technology), we generated a series of 200 bp “packets” of information, which tile the encoded DNA with a pre-defined amount of overlap between each two successive packets (shown in Figure 11 for a 175 bp overlap). These packets can be synthesized as a mixed pool, stored, and sequenced using standard highthroughput sequencing technology, which would allow assembly of the full-length DNA sequence using only the overlaps, without necessitating the complicating addition of addresses and addressing schemes. In order to ensure even coverage at the termini, we generated successively shorter fragments at these areas.

**Figure 11:**
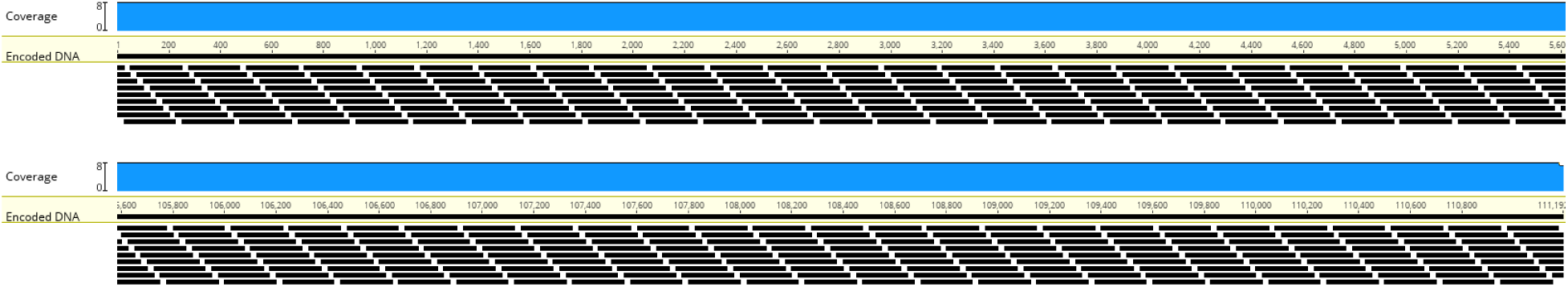
Distribution of oligos along the encoded DNA. Black bars indicate 200 bp oligos, produced so as to tile the encoded sequence with 175 bp overlaps between two successive oligos. Blue graph shows coverage of the DNA by oligos (uniformly 8x virtually everywhere). Only the beginning and end of the sequence is shown here; but the parts shown here are representative of the entire sequence.

#### Simulated sequencing and assembly

After generating packets with varying overlaps (75, 100, 125, 150, 175 bp corresponding to mean sampling densities ranging from 1.6 to 8) we simulated the outcome of sequencing this pool of oligos using the ART sequencing simulator (details in Table 5) at varying read depths (1x, 2x, 5x, 10x, 50x). Afterwards we attempted *de novo* assembly of the resulting FASTQ files, compared the longest resulting contig to the original sequence, and considered two key measures of sequence similarity: How much of the original sequence was present in the resulting contig (recall), and how much of the resulting contig matched the original sequence (precision).

**Table 5:** ART simulator parameters

| **Parameter** | **Value** | **Explanation** |
| --- | --- | --- |
| Sequencer | HiSeq 2500 | Sequencing equipment to be simulated; we chose a commonly used Illumina sequencer |
| Read length | 150 | Length of simulated reads |
| Read depth | 1, 2, 5, 10, 50 | How many reads to collect for a given position. Higher numbers represent deeper sequencing (with more reads covering the same sequence). |

Our measures of precision and recall correlated very closely with each other: With read depths of 5x or more, assembly could easily succeed even when overlaps between successive packets are small. Below this threshold, only comparatively dense tilings of packets could be assembled: With 8x sampling density, even 1x read density was sufficient to recover the original sequence. However, lower sampling densities performed very poorly with low read depth, and contigs assembled would be a fraction of the size of the original sequence, as well as having little similarity to it (indicating spurious assembly becoming the dominant process). A visualization of these findings is provided in Figure 12. Based on these results, we decided to use a sample depth of 8x (175 bp overlap) and a read depth of 10x for subsequent work.

**Figure 12:**
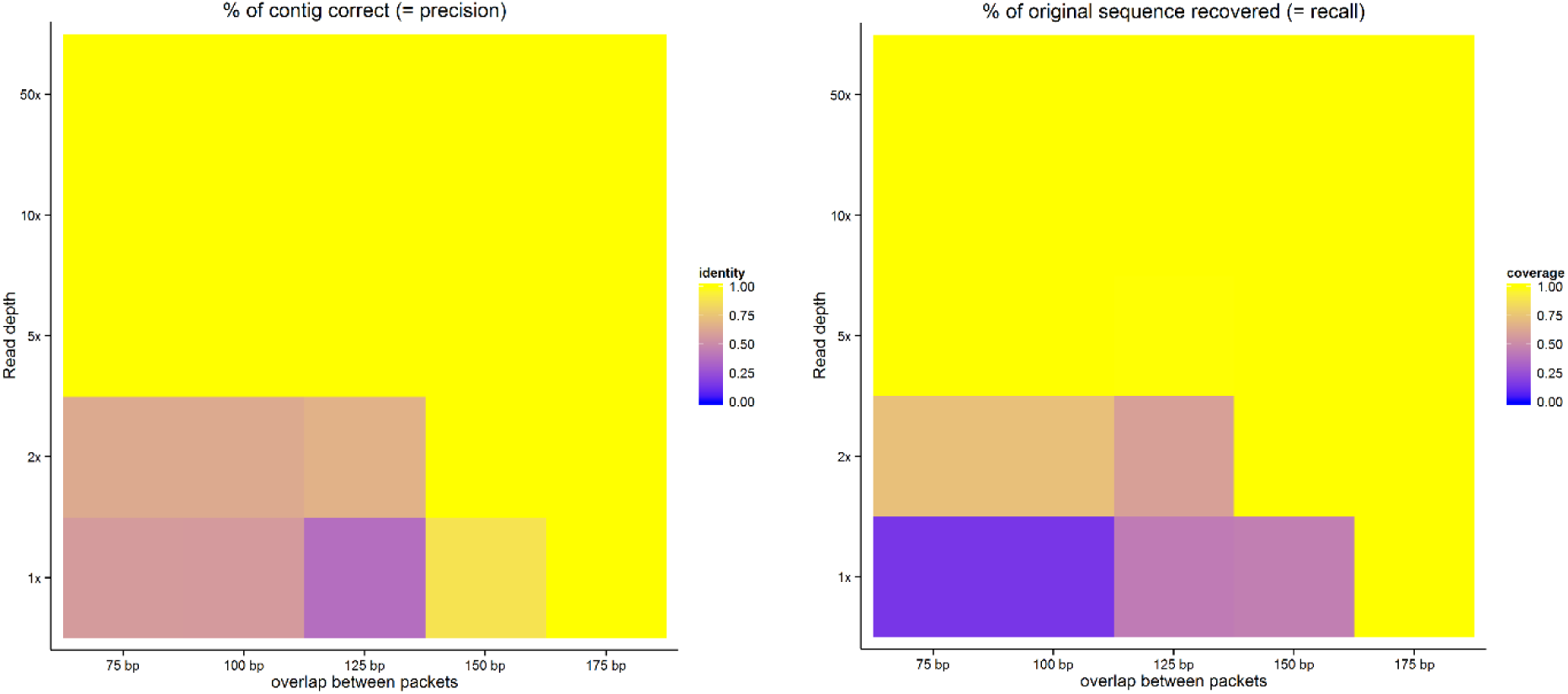
Likelihood of successful assembly at varying read and sampling depths. Left: Fraction of base pairs in the longest assembled contig that matched the original sequence after mapping. Right: Fraction of the original sequence that was present in the longest assembled contig.

Notably, we simulated sequencing of a complex pool of short oligos. Simulated reads obtained from this pool were assembled naively; we did not attempt to recover individual packets, and then assemble packets. Rather, due to the overlaps between successive packets, our assembler was able to seamlessly combine these reads into a single contig without additional intervention. We consider this to be an advantage of our conservative overlap-based segmentation of data.

#### Library construction and long-term packet-wise retention

Because the intended use of our strategy involves cloning the pool of synthesized DNA oligos, there is a risk that not every sequence in the pool will be represented. If too many packets “drop out”, our assembly method may suffer catastrophic failure (a full contig will be impossible to build if a critical number of adjacent packets are missing entirely at one or more locations). However, with higher sampling densities, it is possible that even though one packet is lost during cloning, the two packets adjacent to it will bridge the gap and nevertheless rescue successful assembly. In order to investigate the validity of this tradeoff in the practical context of our work, we performed Monte Carlo simulations of library coverage under the assumption of a defined number of clones being harvested (set to a constant multiple of the sequence diversity).

In all, we conducted 20 simulations each from pools of 100 and 500 unique sequences and expected mean library sample rates ranging from 1 to 30 clones harvested per unique oligo sequence. We then looked at what fraction of the initial set was represented in the draw (Figure 13). As expected, if the total number of clones harvested is equal to the number unique sequences, some sequences appear repeatedly and many are not captured at all. In our case, about a third of the pool would be lost under this cloning regimen. Collecting a very large number of clones virtually guaranteed that no sequence would be lost. Interestingly, the threshold of full coverage was between 10 and 5 clones per unique oligo (cpo): The vast majority of the sequences could be recovered with 5 cpo, but in most simulated runs there would be a few packets missing. On the other hand, if 10 clones per oligo are considered then every single packet was recovered in all 20 simulation runs.

**Figure 13:**
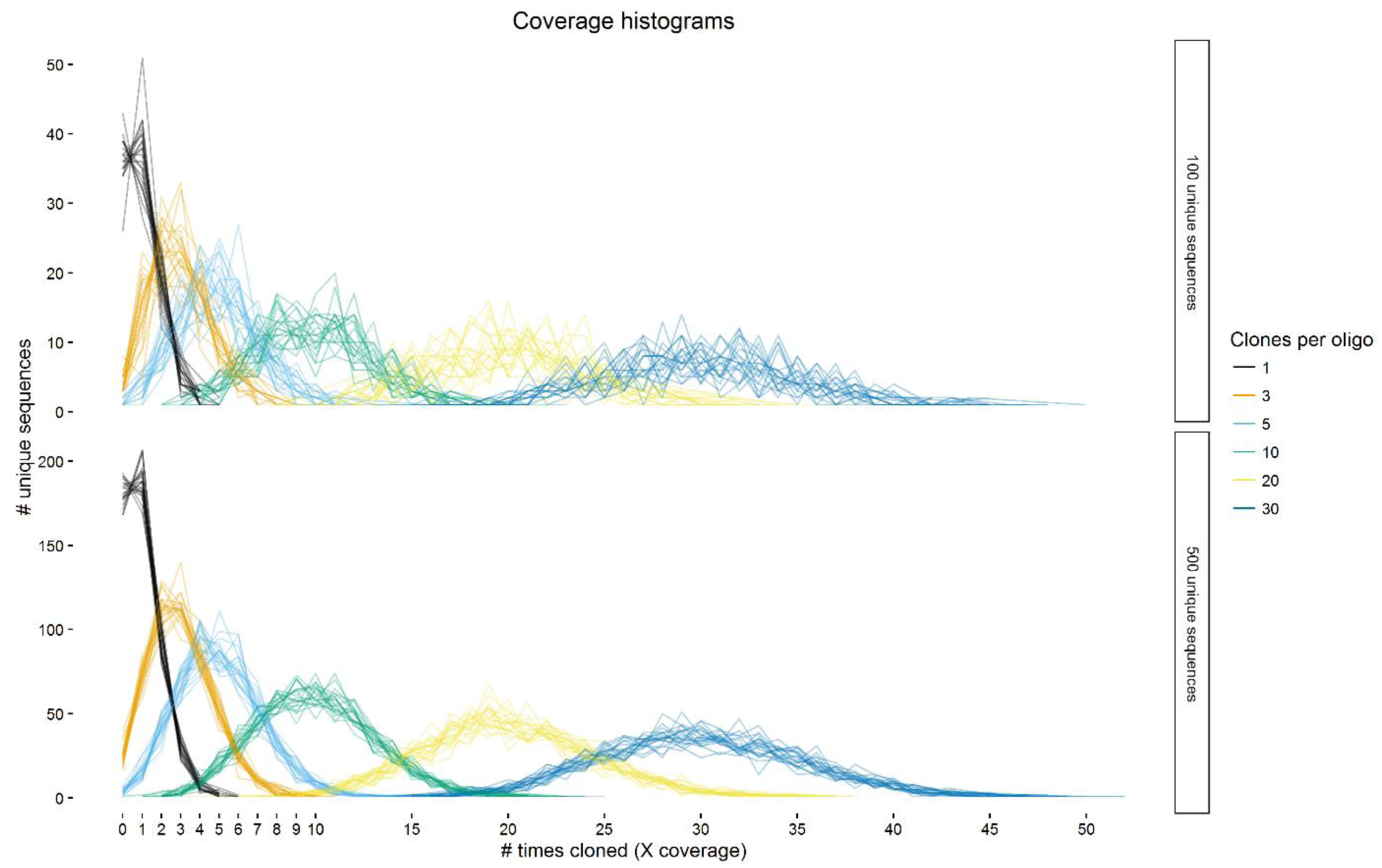
Monte Carlo simulations of library construction from pools of oligos. A series of simulated random draw experiments were performed for pools of 100 and 500 unique sequences; the number of draws ranged from 1 to 30 times the number of unique sequences. For each combination of parameters, 20 repeat experiments were performed, shown here as lines of identical color.

Interestingly, the complexity of the pool *per se* does not appear to have an effect on these boundary conditions. The most pronounced difference between the 100 oligo and 500 oligo case was that the variation was greater with a less complex pool, while the more complex pool behaved more predictably in individual runs.

We concluded, based on our observations, that harvesting 10 clones per oligo is sufficient to have a >95% confidence that all original packets will be well represented.

#### Simulated recovery of information with packet loss

Having observed that harvesting roughly 10 clones for each unique oligo should almost guarantee full library coverage, we attempted to test this inference with *in silico* experiments. We conducted a series of simulated experiments in which the initial pool of oligos was randomly sampled with replacement (since the number of molecules in each class is typically much higher than the number of distinct sequences in synthesized oligo pools, we regarded the effect of replacement as negligible). These random subsamples were then subjected to simulated sequencing with ART at 10x read depth, and then *de novo* contig assembly of the resulting reads was attempted to determine whether recovery of the original sequence is possible even with lost packets. This was repeated 20 times to account for the influence of chance events on recovery.

As shown in Figure 14, results were in line with the expectations arising from the library-coverage experiments. We saw that with a single clone harvested per oligo, in all 20 cases the resulting contigs do not match the original sequence. Moreover, we have seen that due to very poor coverage of the original pool, under this regimen the assembly suffers catastrophic collapse: The resulting contigs do not exceed roughly a third of the original sequence by length, and their sequence often diverges from the reference. Predictably, applying the error correction did not ameliorate this situation.

**Figure 14:**
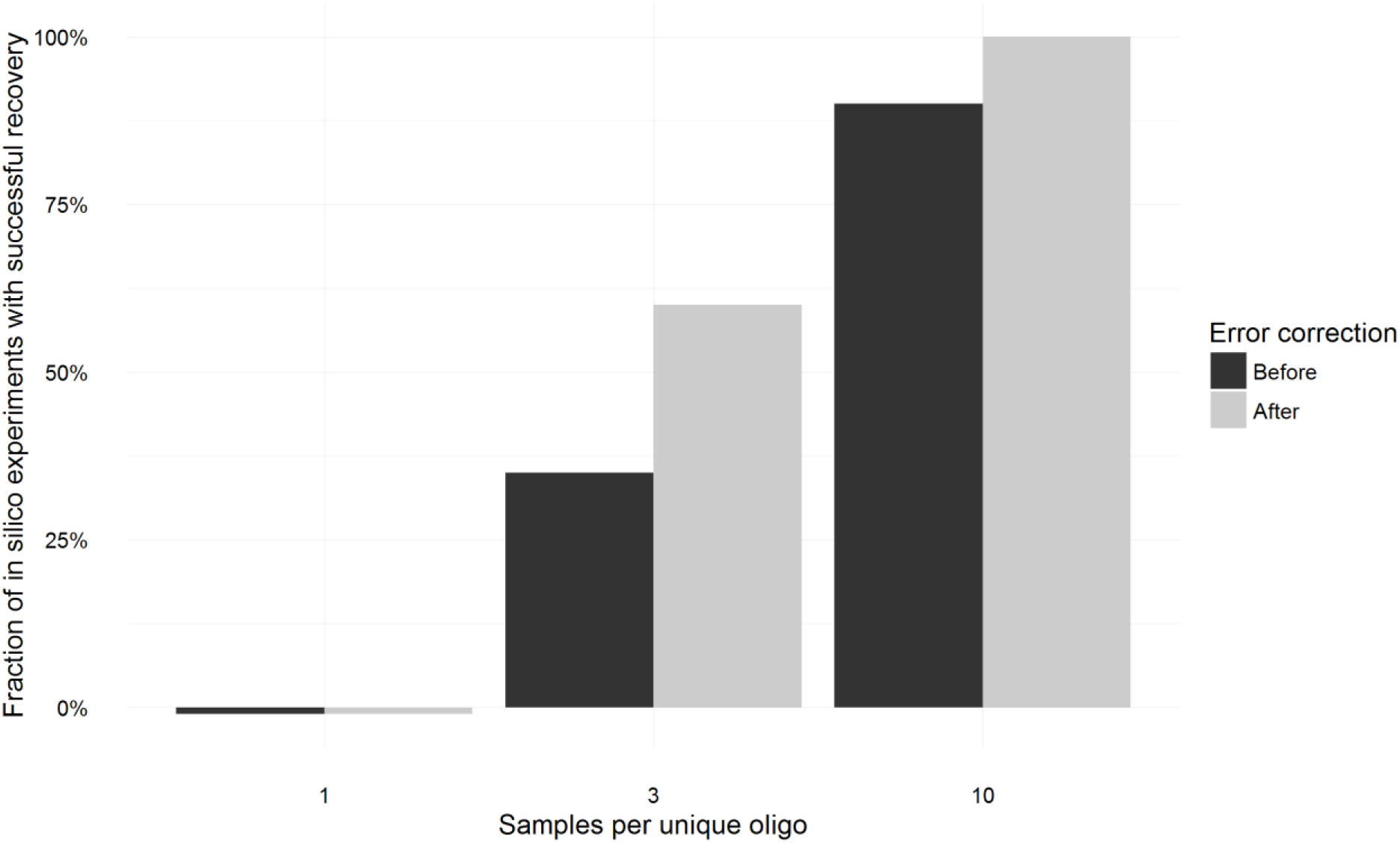
Recovery of original sequence after simulated sampling of packet pool and simulated sequencing. Each bar shows fraction of 20 simulated read-write experiments in which the data after decoding matched the encoded data exactly. For black bars, the error correction capacity built into the sequence was ignored, and the assembled contig was decoded as-is. For grey bars, the error correction was applied and decoding was attempted after.

With the previously established “safe” regimen of 10 clones per oligo, the vast majority of the 20 experiments yielded contigs that were exactly identical to the original sequence, demonstrating successful round-trip read-write of digital data as DNA. In two cases, the assembled contig did not match the original exactly, but the only difference in either case was a single missing terminal nucleotide. Because our error correction method is based on Levenshtein distances, which take into account not only substitution but also deletions, it was possible to correct these problems and restore the original sequence exactly. Thus, considering error correction as well, with a regimen of 10 clones per oligo we could reconstruct the original sequence exactly in every single one of our 20 experiments.

Lastly, of interest was the borderline case of 3 clones per oligo. As expected, this regimen was occasionally able to produce exactly matching sequence, but often the contigs would differ slightly due to missing one terminal nucleotide. Many such errors were readily amendable with the error correction, such that attempting to correct errors produced a dramatic improvement in how many assembled contigs matched the original sequence exactly.

## Discussion

DNA has great potential as a medium of information storage. Indeed, it has been used for this purpose by all living organisms for millions of years. Molecules of DNA are much smaller than digital devices, can be easily copied using both natural and artificial systems and they can be stably maintained for a very long time: The half life of DNA in solution, depending on pH, temperature, and length, can range from a few years to hundreds of thousands of years [9]. In nature, DNA in fossilized bone has been shown to have a half-life as long as 500 years, implying recovery should be possible after many thousands of years given sufficient initial copies of the DNA [5], and useful sequence has been recovered from 450-800 kyr old samples encased in ice [10]. In contrast, digital media degrades quickly over decades [11] [12] [13] [14].

Besides durability of the medium itself, DNA enjoys a unique advantage in that the characteristics of DNA are fixed over time. In contrast, electronic formats change frequently and require specific read/write equipment, technology that can become lost on more epochal time scales [11] [15]. This molecular conservatism will help drive recovery irrespective of what version of DNA sequencing technology is available even in the far future. Taken together, these features highlight the possibilities for DNA for extraordinarily long term archiving.

We have described the design and *in silico* implementation of a strategy for encoding digital information into distributed DNA sequences and then reassembling the original information irrespective of intervening errors. We evaluated several key parameters, including the manner in which the sequence would be partitioned into oligonucleotides, the cloning regimen necessary to maintain a sufficient fraction of the original information for reconstruction following dispersal, the read depth needed to ensure successful recovery, and limits on how many mutations can be tolerated while still allowing complete recovery of the information. Our simulations demonstrate a successful round-trip of information during long term storage in DNA, taking into account problems that might be encountered during synthesis (the write phase) and sequencing (the read phase).

## Useful properties of our encoding strategy

The error correction method we have chosen, and the closely related algorithm for codebook generation, facilitate tailoring the encoding to specific applications. By reducing the minimum distance between code words, one can reduce redundancy, thereby allowing the encoded data to fit into shorter DNA. Conversely, when the integrity of the data is more important than the efficiency of storage, the minimum distance can be set very high, thereby potentially allowing recovery even if a large number of mutations are introduced.

By compressing our input data with LZMA, we remove correlations between the statistical properties of the input data and those of the resulting DNA sequence. LZMA is a compression algorithm based on constructing a Markov model of the data that captures recurring patterns within it, and then encoding the overall data in terms of this Markov model. To the extent that patterns do exist, the resulting information is compressed and largely free of patterns, resembling random data or uniform digital noise. Thus, LZMA should allow us to produce uniform and non-repetitive DNA sequence from even very skewed and / or repetitive inputs. By explicitly generating a codebook with well-defined parameters, we should be able to further impose constraints on any resulting sequence output, such as requiring a GC composition that would be optimal for synthesis and sequencing, irrespective of the composition or structure of the input data.

In our work we used a codebook of eight 4 bp blocks. In principle, it is possible to obtain the same level of overall redundancy with longer blocks, the minimum codeword distance being scaled up as appropriate. Two key parameters for choosing the block size were performance (codewords for longer blocks take more time to generate) and the expected distribution of mutations. Longer codewords, with more distance between them, have a greater per block mutation tolerance, even if the per base pair mutation tolerance remains the same. Error correction functions only if the number of mutations within a particular block remains below a threshold; that is, mutations must not be too close to each other. If there is a tendency for mutations to occur in small clusters, rather than being evenly distributed, longer blocks would likely be preferable.

## Relation to previously published literature

DNA-based information storage has previously been explored, as challenges with conventional digital storage methods became apparent. Bancroft *et al.* 2001 provided an early proof of concept and laid out the theoretical framework for encoding data into DNA. They proposed an encoding scheme (which they consider “obvious”) that maps triplet combinations of the three nucleotides A, C, T to ternary numbers and uppercase letters of the English alphabet, along with a space character. The information is subdivided into segments (called iDNAs), and each segment is prefixed with a spacer and unique primer sequence, and then flanked with universal forward and reverse primers. An ordered array of unique primers is also included as a “polyprimer key”, also flanked by the universal primer. With this, they encode the famous opening of *A Tale of Two Cities* by Charles Dickens in two iDNAs 232 and 247 bp long. The iDNAs can then be individually recovered by PCR and sequenced. Notably, this conception predates the revolution in reading DNA sequences wrought by NextGen sequencing technology, which we extensively leverage in our own work.

Church and colleagues reported in 2012 the storage and subsequent recovery of a 5.27 megabit stream of information (composed of text, images and source code). The information was encoded with a degenerate code, mapping A or C to 0 and G or T to 1 (this provided some freedom in preventing homopolymer runs and GC-skew). The information is partitioned into 54,898 blocks 159 bp each, of which only 96 bp encodes information while 19 nt is used for address blocks and 44 bp for universal primers (incidentally this implies a rate of about 0.3 input bits per output bit, very close to that which we calculate for our encoding strategy). After sequencing and filtering for perfect reads, they obtain 3000-fold coverage of each piece (for comparison, our simulations demonstrate data recovery with 5-fold and 10-fold sequencing). However, their demonstration still contained 22 nucleotide errors, of which 10 were also bit errors. Interestingly, this work notes that future work could incorporate compression, redundant encodings, parity checks and error correction - three of which are central aspects of our own work.

Goldman 2013 describes the encoding and reconstruction of various computer files totalling 739 kb. The digital information was first Huffman coded (which reduces the space taken up by frequently occurring symbols) and then converted to a ternary representation, which is encoded into DNA at one nucleotide per ternary digit, rotating the nucleotides to prevent sequences that are difficult to synthesize or sequence from occurring. This single stream of DNA is broken into 100 bp fragments with 75 bp overlaps similar to our approach, but that alternate forward and reverse complement information in subsequent pieces. The overlaps are used only for redundancy, not assembly, and instead an address block is added to each piece to enable assembly. Their final encoding produces 153,335 DNA sequences that are each 117 bp long, including the address information, which corresponds to a rate of roughly 0.67 (ignoring the four-fold redundancy from overlaps). After sequencing and filtering, they obtained a set of reads with 1308x coverage, from which they were able to recover the encoded digital data after applying error correction. For comparison, in our case we simulated read depths of 5x and 10x, after which all of the data was recovered perfectly without need for error correction, with the exception of a few cases in which a single terminal nucleotide was lost. These losses could nonetheless be rectified by applying error correction so that the original data was recovered intact.

Our strategy represents several important innovations over previous work. Most importantly, we do not rely on address blocks to guide assembly of the DNA sequence for decoding, but instead make use of the overlaps between packets of data. These overlaps can be used to construct a De Bruijn graph and be assembled in a manner analogous to the well-studied problem of genome sequencing and assembly. A primary advantage of this method is that only standard assembly algorithms, not specialized software, are required to perform the assembly. In addition, the different parts of the packet are equally vulnerable to mutation. With an address scheme, mutations falling on the address section can lead to the loss of the whole block, while mutations falling on the data portion are far more limited in scope. In our approach, if the packets tile the original sequence uniformly, it does not matter where the mutations fall, provided they are sufficiently sparse. This greatly simplifies the design of a single universal error correction scheme. Finally, the overlaps act as additional mutation buffers. Our encoding has tunable redundancy that already confers an error-correction capacity, but the added redundancy from overlaps allows straightforward detection and correction of sequencing errors and rare mutations as a result of the de novo assembly process. This ‘failsafe’ error correction protocol makes this strategy the most secure thus far for any long term storage attempts.

With address blocks, an important consideration is efficiency of storage: within each packet, the address information and the encoded data compete for space. The number of packets scales with the total size of the encoded digital data, and the associated address block becomes larger when there are ever more packets. An upper bound on the number of packets is the number of possible unique sequences of that length (in reality this tends to be a gross underestimation since many possible sequences are unsuitable for synthesis and sequencing). Goldman and colleagues note, for instance, that with 114 bp packets, 14 bp could be used for indexing to obtain 88% efficiency; this imposes an upper limit of about 268 million packets covering 6.7 Gbp of sequence (taking into account the four-fold redundancy). For a similar level of redundancy, our approach could generate 114 bp packets tiling the sequence with 85 bp overlaps, for a total of 231 million packets. The upper limit on how much information can be stored would most likely be determined by the limits of assembly software. Moreover, arbitrarily large amounts of information can be stored regardless of this upper limit, so long as encoded DNA pools are somehow stored and sequenced separately.

Our overlap-based approach therefore allows for a simpler and more straightforward partitioning of encoded data than relying on address blocks. It eliminates the need for parsing and interpreting address blocks when decoding, as well as the concern over how to preserve the instructions for performing this step correctly. The sizes of address blocks must be explicitly standardized prior to encoding, which will impose an upper limit on how much data can be encoded for as long as that standard is in effect. If the address blocks are too small, large encodings will run out of address space and the encoding standard will have to be changed often (leading to compatibility issues), but if they are too big then small pieces of data will be encoded inefficiently. Our approach sidesteps this dilemma; it is only necessary that sufficient overlap exists between packets to enable assembly, and no space is “wasted” on address blocks since the overlaps act as an additional safeguard of data integrity. The information is partitioned efficiently regardless of its size, the upper bound on capacity is very large, and no specific details of the partitioning need be recorded, since simply attempting to assemble the overlapping pieces of DNA (a step which would be obvious even if the encoding scheme is unknown) yields the complete stream of encoded data.

## Applications and interdisciplinary context

Though our primary interest was to develop the means of storing information in DNA, the process can be adapted to a number of other applications. Herein we will discuss four of these: repurposing the codebook generator to produce barcodes, using encoding for inactivation of toxic sequences, the potential to use DNA-encoding to protect private information, and bridging extinction events.

We designed our codebook generation algorithm to produce a set of sequences that are evenly spaced in sequence space. This encoding has similarities to the mathematical concept of a ‘perfect code’ (MacWilliams 1977) which refers to sets of codewords optimally arranged in sequence space so as to provide a given level of error correction, although our approach considers Levenshtein distances rather than Hamming distances, and furthermore there are codewords we deliberately avoid, so in practice our code is less than perfect. Though our purpose was to encode information, an interesting property of the resulting codewords is to lack bias and repetition, and therefore be distinct from one another. The relatively small length makes them easy to synthesize and the lack of repetition makes them easy to sequence. Overall, these properties also potentially make them excellent DNA barcodes [16] [17] [18], and as such they can be readily applied to a long list of biological techniques such as multiplexed NextGen sequencing [19], identification of genetic variants [20], construction of deletion libraries [21] and high-throughput RNA profiling [22].

Another consequence of encoding is that output sequence has very little resemblance to input. This can make the storage of difficult sequences more tractable. For example, if a given DNA sequence is fragile or unusually prone to mutation, encoding should allow it to be stored with high fidelity. Our centromere simulations illustrate this principle - although centromeric sequence is not *per se* toxic, it is very repetitive, impairing its faithful construction and sequencing. In contrast, when encoded, centromeres do not have any obvious repeats or other unwieldy sequences. Encoding can be used to not only preserve information, but also to prevent its ready expression in the absence of decryption. Any biological sequence is rendered biologically unreadable by our scheme for encoding, and this in turn suggests that biothreats such as toxin genes or even entire pathogen genomes (e.g. genomes of infectious viruses) that would otherwise be harmful in their ‘biologically readable’ form could potentially be encoded and stored over long periods, essentially placing them in ‘deep storage’ for posterity.

Distinctly from previous work, we encode information through an explicit codebook, which then becomes necessary for later decoding. In this sense, our encoding method is analogous to a symmetric key cryptography scheme, with the codebook as the key. With short block lengths, it is conceivable that the codebook might be reverse engineered from the encoded sequence only by an exhaustive approach. With longer block lengths, it would become extremely difficult to recover the information without access to the codebook. This allows information to be hidden from readers who do not posses the correct codebook, or simply to prevent information from being accessed before an arbitrary amount of time has passed (that is, the time needed for exhaustive reconstruction of the codebook). These features render DNA encoding inaccessible and resilient in a way that far exceeds what can be availed via electronic encoding. While there may be few applications for information privacy that can tolerate slower read-write capabilities, it should be noted that mechanical, Enigma-like rotor cypher systems can be difficult and time-consuming to crack even with modern electronic computers [23]. Thus, it is entirely possible that a nation state or corporate enterprise might wish to hide (and retrieve on a leisurely scale) ‘trade secrets’ *via* the long term, difficult to break, and undegradable encoding strategies we describe herein.

Finally, because of the Markov chain-based compression employed, having access to part of the encoded DNA does not allow one to access part of the original information. Decoding is only practical if the entire sequence is present, since otherwise both assembly and decompression fail. This all-or-nothing nature of the information round trip requires care to be taken that a sufficient number of packets are retained throughout storage; in our case we solve this by creating a redundant set of packets that allow assembly even if a small number is lost. Subsets of the packets could also be distributed to different recipients or locales, which would then be unable to decode the information on their own without combining their respective pieces of the DNA-encoded data. This strong message compartmentalization, combined with the potential for multi-hundred thousand year digital file storage capacities–a time-scale exceeding all known continuous human civilizations–allows one to imagine even more fantastic scenarios: For example, the distribution of DNA encoded information and the relevant codebooks on a large scale might provide a means to reboot the information infrastructure of an advanced society irrespective of the apocalyptic calamities that might befall it. The requirement of retaining or reinventing DNA sequencing technologies, coupled with the need to reacquire a full set of information packets would even lead to situations in which information recovery was only possible once some degree of technological and social reorganization had occurred, post-Apocalypse.

To summarize, we have demonstrated a strategy for encoding digital information into DNA, which is highly parallelizable and has built-in error correction ability, and relies on an overlap-based assembly method fundamentally different from the previously published approaches. We have demonstrated full round trip read and write of the information, including simulated sequencing of a cloned oligo pool, with recovery from simulated mutations.

## Acknowledgements

We are immensely grateful to Randall Hughes for invaluable discussions regarding the technical details of DNA synthesis technology and its limitations. E.M.M. acknowledges grant support from the NIH, NSF, CPRIT, and Welch Foundation (F-1515).

